# DNA damage models phenotypes of β cell senescence in Type 1 Diabetes

**DOI:** 10.1101/2021.06.07.447428

**Authors:** Gabriel Brawerman, Jasmine Pipella, Peter J. Thompson

## Abstract

**Objective:** Type 1 Diabetes (T1D) is characterized by progressive loss of insulin-producing pancreatic β cells as a result of autoimmune destruction. In addition to β cell death, recent work has shown that subpopulations of β cells acquire dysfunction during T1D. We previously reported that some β cells adopt a senescent fate involving a DNA damage response (DDR) during the pathogenesis of T1D, however, the question of how senescence develops in β cells has not been investigated.

**Methods:** Here, we tested the hypothesis that unrepaired DNA damage triggers β cell senescence using culture models including the mouse NIT1 β cell line derived from the T1D-susceptible nonobese diabetic (NOD) strain, human donor islets and EndoC β cells. DNA damage was chemically induced using etoposide or bleomycin and cells or islets were analyzed by a combination of molecular assays for senescence phenotypes including Western blotting, qRT-PCR, Luminex assays, flow cytometry and histochemical staining. RNA-seq was carried out to profile global transcriptomic changes in human islets undergoing DDR and senescence. Insulin ELISAs were used to quantify glucose stimulated insulin secretion from chemically-induced senescent islets and cells in culture.

**Results:** Sub-lethal DNA damage in NIT1 cells led to several classical hallmarks of senescence including sustained DDR activation, growth arrest, enlarged flattened morphology and a senescence-associated secretory phenotype (SASP) resembling what occurs in primary β cells during T1D in NOD mice. Some of these phenotypes differed between NIT1 cells and the MIN6 β cell line derived from a non-T1D susceptible mouse strain. RNA-seq analysis of human islets undergoing DDR and early senescence revealed a coordinated p53-p21 transcriptional program and upregulation of prosurvival signaling and SASP genes, which were confirmed in independent islet preparations at the protein level. Importantly, chemically induced DNA damage also led to DDR activation and senescence phenotypes in the EndoC-βH5 human β cell line, confirming that this response can occur directly in human β cells. Finally, DNA damage and senescence in both mouse β cell lines and human islets led to decreased insulin content.

**Conclusions:** Taken together, these findings suggest that some of the phenotypes of β cell senescence during T1D can be modeled by chemically induced DNA damage in mouse β cell lines and human islets and β cells in culture. These culture models will be useful tools to understand some of the mechanisms of β cell senescence in T1D.

**Highlights:**

- DNA damage induces senescent phenotypes in mouse β cell lines
- DNA damage induces a p53-p21 transcriptional program and senescent phenotypes in human islets and EndoC cells
- DNA damage and senescence leads to decreased insulin content
- DNA damage models some aspects of β cell senescence in Type 1 Diabetes

## 1. Introduction

Type 1 Diabetes (T1D) is a disease of chronic insulin deficiency due to progressive autoimmune-mediated destruction of pancreatic β cells [1]. While much of our understanding of T1D stems from investigations of the immune system, recent work has implicated stress responses in β cells as an additional factor driving disease progression and onset [2, 3]. For instance, β cells experience unmitigated endoplasmic reticulum (ER) stress and activate the terminal unfolded protein response (UPR) leading to apoptosis during the development of T1D [4]. Small molecule therapeutics that block terminal UPR have shown promise in clinical trials of new onset T1D patients by slowing the decline in C-peptide production after diagnosis [5, 6]. Thus, clinical interventions to target β cell stress responses are emerging as a therapeutic strategy for T1D.

In addition to terminal UPR, it was recently shown that a subpopulation of β cells undergo senescence during the pathogenesis of T1D [7]. Cellular senescence is a complex and dynamic program of irreversible cell cycle arrest, typically initiated as a consequence of aging, oncogene activation, or a variety of stressors, and has many different biological functions including tumor suppression, wound healing, tissue regeneration and embryonic patterning [8, 9]. Senescence resulting from unrepairable cell stress or damage is usually activated via a combination of the p16^Ink4a^/Rb and/or p53-p21 pathways, leading to permanent growth arrest and hallmark phenotypes such as an enlarged cell morphology, senescence-associated βgalactosidase (SA-βgal) activity, persistent DNA damage response (DDR), apoptosis resistance and a senescence-associated secretory phenotype (SASP) [10].

While healthy β cells are progressively destroyed by autoimmunity in T1D, senescent β cells accumulate during the development of T1D in humans and in the preclinical nonobese diabetic (NOD) mouse model for T1D [7]. Senescent β cells in NOD mice are characterized by upregulation of p21 and p16^Ink4a^, DDR activation, a Bcl-2-mediated prosurvival phenotype, elevated SA-βgal activity and a SASP involving IL-6, Igfbp3 Serpine1, Mmp2, and Flnb [7]. The presence of persistent DDR, Bcl-2 upregulation and contents of SASP generally differentiates stress-induced senescence in β cells in T1D from natural aging and β cell senescence in Type 2 Diabetes [11–13]. Notably, in NOD mice, elimination of senescent β cells with Bcl-2 family inhibitors (senolytic compounds), or suppression of SASP with Bromodomain ExtraTerminal domain inhibitors, halts disease progression [7, 14]. Therefore, senescence is a novel therapeutic target for T1D prevention.

Despite these advances, basic mechanistic questions around β cell senescence in T1D remain. Senescence progression and phenotypes are variable between cells and are generally cell type-and stress/trigger-dependent [15, 16] and although well studied in vitro, are poorly understood in vivo, as there is no one marker that can accurately define senescence in every cell type [17]. However, culture models have been foundational in defining and characterizing molecular mechanisms and phenotypes. In particular, the specific triggers of β cell senescence in T1D are not known and there is currently no β cell culture model to study the phenomenon. Different inbred mouse strains used to model diabetes have different genetic background polymorphisms that could influence the development of specific senescence phenotypes. For instance, the NOD mouse strain commonly used to model T1D harbors two polymorphisms in the non-homologous end joining repair factor Xrcc4, lowering the efficiency of DNA double-strand break repair in this strain as compared with C57BL strains [18]. Addressing fundamental mechanistic questions around β cell senescence in T1D using mouse models would greatly benefit from a cell line with the relevant genetic background.

As with NOD mice, a subset of β cells in human T1D donors show DDR and activation of the p53-p21 senescence pathway [7, 19]. We previously showed that cultured primary adult human islets transcriptionally upregulate p21 (*CDKN1A)* and develop a SASP following induction of DNA double-strand breaks with the chemotherapeutic agent bleomycin [7, 14]. These findings suggest that β cell senescence in T1D may involve unrepaired DNA damage, but a comprehensive analysis of the outcomes of DDR induction in human islets has not been performed and it is unclear whether these changes actually occur in β cells, as islets are comprised of a diversity of cell types.

The objectives of this study were to investigate the phenotypes arising in β cells as a result of DDR activation to test the hypothesis that sub-lethal DNA double-strand break damage triggers senescence in β cells derived from the NOD mouse strain and in human β cells. Our results suggest that chemically triggered DNA damage can be used to accurately model some of the key phenotypes involved in β cell senescence during T1D. We also found that chemically induced DDR and senescence decreases insulin content in mouse β cell lines and human islets. These findings have important implications for culture modeling of β cell senescence and understanding the mechanisms driving senescence in the context of T1D.

## 2. Materials and Methods

### 2.1 Mouse cell line culture and drug treatments

Mouse insulinoma cell lines MIN6 (RRID:CVCL_0431, sex of cell line is unknown) derived from 13 week C57BL6 mice (gift from Dr. A. Bhushan, UCSF) and NIT1 (RRID: CVCL_3561), derived from 10 week female NOD/LtJ mice (purchased from ATCC) have been previously described [20, 21]. Both cell lines were cultured in DMEM high glucose with 2 mM L-glutamine (ThermoFisher) containing 10% FBS (Millipore-Sigma), 20 mM HEPES, 1 mM sodium pyruvate (Gibco), 1X antibiotic-antimycotic (Gibco) and 50 µM 2-mercaptoethanol at 37°C and 5% CO_2_. Cell lines were confirmed mycoplasma-free using the PlasmoTest mycoplasma kit (Invivogen). Cell lines were passaged every 4-6 days and MIN6 were detached with 0.25% trypsin-EDTA (Gibco) and seeded at 7.5-1×10^6^ cells per 75 cm^2^ flask, whereas NIT1 were detached with 0.05% trypsin-EDTA and seeded at 2-2.5×10^6^ cells per flask. Cells were used between passages 4 and 25 for all experiments. For senescence induction, we modified a previous method that was used to trigger “therapy-induced” senescence in mouse melanoma cell line B16-F10 [22]. MIN6 cells were seeded at a density of 0.7-0.85 x 10^5^ cells/cm^2^ in 12 or 6-well plates, respectively. Twenty-four hours later, normal growth media was replaced with media containing 2 µM etoposide (Millipore-Sigma) or vehicle (0.01% DMSO) for 2 or 3 days, followed by removal of drug by replacing wells with fresh media. Similarly, NIT1 cells were seeded at a density of 0.85-1.15 x 10^5^ cells/cm^2^ and 24 h later cultured in 0.5 or 0.25 µM etoposide or vehicle (0.0025 or 0.00125% DMSO) for 3 days before drug removal on day 4 as with MIN6 cells. After drug removal, the media was replaced with fresh media every other day. Vehicle-treated cells often reached confluence before drug washout and required passaging prior to drug removal, so cells were passaged into the same concentration of DMSO-containing media and media was replaced at the same time as the etoposide-treated cells. Cell counting and viability assays were performed with trypan blue staining on an automated cell counter (Bio-Rad TC-20). Cells were cultured for indicated times after drug treatment and harvested as indicated for various assays.

### 2.2 Human islet culture and EndoC-βH5 cells

Human islets were procured either through the Integrated Islet Distribution Program (IIDP) (for RNA-seq) or from the Alberta Diabetes Institute (ADI). Islets used for RNA-seq were isolated from a healthy nondiabetic female donor of Hispanic ethnicity, age 44 years (RRID:SAMN13186657), from IIDP. Islets from ADI were used for DDR induction and Luminex assays were from a healthy nondiabetic Caucasian male donor age 32 years (RRID:SAMN 22441940). Islets from ADI were used for DDR induction, GSIS and western blots were from a healthy nondiabetic Caucasian male donor age 55 (RRID: SAMN23408101). Upon receipt islets were rested in culture for 24 h in RPMI-1640 media containing 10% FBS, 2 mM L-glutamine, 5.5 mM glucose and 1X antibiotic-antimycotic, as previously [7, 14]. To induce DDR and senescence, islets were cultured in media containing 50 µM bleomycin (Medchem express) or vehicle (0.2% DMSO) controls for 48 h, followed by removal and culture in drug-free media for 2-5 days, as previously [7]. Islets were then harvested for RNA extraction or protein. For conditioned media, islets were put into serum-free islet media on day 4 post-drug washout, for 24 h and the conditioned media collected. To induce ER stress and UPR, islets were cultured in 1 µM Thapsigargin for 5 h, before harvest for protein extraction.

The human fetal EndoC β cell lines have been previously described [23]. The latest derivative, EndoC-βH5, were purchased from Human Cell Design. These cells are derived from the earlier EndoC-βH3 line [24] and have clonally selected for maximal insulin secretory responses and β cell identity marker expression (Human Cell Design). The βH5 line has undergone tamoxifen-mediated excision of the immortalizing SV40 Large T-antigen and hTERT transgenes as in the earlier EndoC-βH3 cell line to render the cells mature and non-proliferative. βH5 cells were seeded at 3.75 or 4×10^5^ cells per well in 12-well TPP plates treated with gel matrix provided by the supplier (βCOAT) and cultured in serum free media (OPTIβ1) as previously, containing: DMEM that contained 5.6 mM glucose, 2% BSA fraction V, 50 μM 2-mercaptoethanol, 10 mM nicotinamide, 5.5 μg/ml transferrin, 6.7 ng/ml selenite, 100 U/ml penicillin, and 100 μg/ml streptomycin (supplied by Human Cell Design). After thawing cells, media was changed within 4 h after cells attached and was replaced with media containing 0.1% DMSO (vehicle control) or 35 μM bleomycin for 48 h. Following drug treatment, the media was replaced with fresh media for an additional 2 days of culture or cells were harvested immediately after drug treatment for western blot. In a second experiment cells were cultured in normal growth media for an additional 4 days after drug removal and media was replaced on day 4 to generate conditioned media for 24 h for Luminex assays

### 2.3 Western blot analysis

Whole cell extracts were prepared with RIPA buffer (ThermoFisher) in the presence of 1X protease and phosphatase inhibitor cocktail (ThermoFisher) and protein was quantified with the BCA assay (ThermoFisher). Testis tissue from C57BL6/J 3 month old mice was a gift from Dr. C. Doucette (University of Manitoba). Approximately 3-10 µg of total protein from testis, MIN6 or NIT1 cells, human islets, or EndoC-βH5 cells treated as indicated (equal amounts per sample for each experiment) were electrophoresed on precast 4-12% Bis-Tris Plus Bolt Mini gradient gels and electroblotted onto nitrocellulose membranes with the Mini Bolt-iBlot2 system (ThermoFisher). Membranes were blocked in 5% non-fat milk in Tris buffered saline with 0.1% Tween-20 (TBST) for 1 h at room temperature. Primary antibodies used for westerns and dilutions used are indicated in **Supplementary Table 3**. Primary antibodies were incubated on membranes in TBST overnight at 4°C, washed three times for 5 minutes in TBST and detected with HRP-conjugated secondaries (Jackson Immunoresearch, diluted 1:50,000-1:100,000) after incubation for 1 h at room temperature in 5% non-fat milk in TBST. Membranes were washed three times for 5 minutes in TBST and were then incubated with SuperSignal West Pico Plus ECL (Thermo Fisher). Membranes were developed after exposure to X-ray film (Fuji Medical) and two or three exposures were collected to ensure signals were not saturated. Densitometry was performed using ImageJ to quantify band intensities relative to β-Actin as a loading control on each blot.

### 2.4 SA-βgal activity assay

SA-βgal activity in MIN6 and NIT1 cells was detected using a commercial kit (Cell Signaling Technology) according to the product instructions. Cells were subsequently washed in PBS and imaged on an EVOS color brightfield imager.

### 2.5 EdU labeling assay

EdU incorporation was measured as an assay for DNA synthesis using a commercial kit (Abcam) according to the product instructions modified for adherent cells. Briefly, MIN6 or NIT1 cells were induced to senescence as indicated above and on the day of drug removal (72 h post-treatment) cells were incubated with media containing 5 µM EdU for 3h at 37°C and 5% CO_2_. Cells were subsequently washed with d-PBS, trypsinized, counted and resuspended at 1-5 x 10^6^ cells/ml in d-PBS. The zombie NIR fixable viability kit (BioLegend) was used according to the manufacturer’s instructions prior to fixation in 4% formaldehyde. After fixation, cell pellets were washed twice using cell staining buffer (BioLegend), permeabilized and treated with EdU detection solution. After two more washes, cells were filtered through a 70 μm strainer (ThermoFisher Scientific) into 5 mL flow tubes (ThermoFisher Scientific). Flow cytometry was performed on an Attune acoustic focusing cytometer with a gating strategy as follows: FSC-A/SSC-A (cells), FSC-A/FSC-W (single cells), Zombie Near IR-negative (RL3, live cells) and EdU-positive (RL1). Approximately 4-10,000 live cells were scored per sample.

### 2.6 Luminex assays

Conditioned media (CM) was collected by culturing cells or islets in growth media lacking FBS for 18-24 h. CM was collected, clarified by centrifugation (3000g for 5 minutes at room temperature) and either used immediately or stored at 4°C for up to 2-3 weeks. Luminex assays were carried out on CM from cells treated as indicated using a custom magnetic bead kit that included mouse SASP factors IL-6, Serpine1 and Igfbp3, or human SASP factors CXCL1, IL-8, IGFBP4, GDF-15, or TNFRSF10C (R&D systems). 50 µl of CM was assayed on a Bio-Plex 200 luminex platform (Bio-Rad) as previously [7, 14] and secreted amounts in 1-2 ml of CM were normalized to viable cell counts to calculate amount secreted per 100,000 cells or for human islets secretion was normalized to total RNA content. Alternatively, media was spin-concentrated using a 3 kDa MWCO centrifugal unit (Millipore-Sigma) and the concentrated conditioned media was analyzed by Luminex.

### 2.7 RNA extraction and quantitative RT-PCR

Total RNA was extracted from MIN6, NIT1, human islets or EndoC-βH5 cells using a commercial column-based purification with DNAse I treatment (Direct-Zol RNA MicroPrep with Trizol Reagent, Zymo Research). RNA was quantified by Nanodrop spectrophotometer (ThermoFisher Scientific). Approximately 50-300 ng of total RNA was used for cDNA synthesis with random primers (LunaScript SuperMix Kit, New England Biolabs), depending on the cells used. qPCR was carried out on a Bio-Rad CFX96 using Luna Universal qPCR Master Mix (New England BioLabs). Primers used for qPCR were previously validated from other studies and are indicated in **Supplementary Table 3**.

### 2.8 RNA-seq on human islets

Approximately 1 µg of total RNA from bleomycin and vehicle-treated islets (n = 3 biological replicates per group) at 4 days post-drug removal was submitted for paired-end RNA sequencing on Illumina Nova-Seq 6000 platform by a commercial service provider, LC Sciences. Total RNA was quality checked and ribosomal RNA depleted. Libraries were prepared with Illumina TruSeq-stranded total RNA sample protocol and quality checked by Agilent BioAnalyzer 2100 High Sensitivity DNA Chip. Cutadapt [25] and perl scripts were used to filter adapter contamination and undetermined bases. Sequences were quality checked with FastQC (https://www.bioinformatics.babraham.ac.uk/projects/fastqc)and mapped to the human genome (version 96) using Bowtie2 [26] and HISAT2 [27]. Mapped reads were assembled for each sample with StringTie [28]. A comprehensive transcriptome was prepared using perl scripts and gffcompare (https://github.com/gpertea/gffcompare/). StringTie and edgeR[29] were used to estimate transcript expression levels. LncRNAs were identified based on filtering transcripts overlapping with known mRNAs, known lncRNAs and transcripts <200 bps. CPC [30] and CNCI [31] were used to predict transcripts coding potential. Transcripts with CPC score <-1 and CNCI score <0 were removed. Remaining transcripts were considered lncRNAs. StringTie was used to perform differential expression level analysis for mRNAs and lncRNAs by calculating FPKM (total exon fragments/mapped reads(millions) x exon length (kb)). A complete list of differentially and all expressed genes is shown in Supplementary Table 1. A complete list of differentially expressed lncRNAs and novel lncRNAs is found in Supplementary Table 2. Differentially expressed mRNAs and lncRNAs were selected with log2 fold-change >1 or <-1 between vehicle control and bleomycin treatment and parametric F-test comparing linear nested models (p < 0.05) by edgeR. GO and KEGG analysis was performed using by hypergeometric distribution fitting (p < 0.05) from gene ontology terms (http://www.geneontology.org) and KEGG pathways (http://www.kegg.jp).

### 2.9 Static glucose stimulated insulin secretion assays

Glucose-stimulated insulin secretion was measured from MIN6 and NIT1 cells and human islets according to standard methods with minor changes [32, 33]. For mouse β cell lines, cells were seeded as indicated above cultured with vehicle or etoposide for 72 h to induce senescence. At the 72 h post-treatment timepoint, cells were washed in and incubated for 1 h in a tissue culture incubator in Krebs-Ringer Bicarbonate (KRB) buffer (128.8 mM NaCl, 4.8 mM KCl, 1.2 mM KH_2_PO_4_, 1.2 mM MgSO_4_, 2.5 mM CaCl_2,_ 5 mM NaHCO_3_, 10 mM HEPES) containing 0.1% BSA fraction V (Millipore-Sigma). Then cells were incubated for 1 h in fresh KRB buffer with BSA as above containing 0 mM glucose as the low glucose condition, followed by 1 h in KRB with BSA containing 20 mM glucose as the high glucose condition. Total insulin content was collected by lysing cells in an acidified ethanol solution (0.15 M HCl, 95% ethanol). Human islets were treated in n = 6 biological replicates of vehicle control (DMSO) or 50 μM bleomycin for 48 h to induce senescence, followed by culture in drug-free media for an additional 4 days. At day 5 post-drug removal, islets were washed in KRB + 0.1% BSA containing 2 mM glucose (low glucose) and incubated in the low glucose buffer for 1 h. Islets were then washed into KRB + 0.1% BSA containing 20 mM glucose for 1 h (high glucose buffer) and supernatants of the 2 mM and 20 mM glucose buffers were collected. For both mouse β cell lines and human islets, aliquots of the low and high glucose supernatants and the insulin content extracts were collected and spun at 3000g for 5 minutes at 4°C and stored at −20°C or used immediately for mouse or human insulin ELISAs (Mercodia) with appropriate dilutions (1:10-1:50 for supernatants, 1:100-1000 for insulin content). Insulin secretion in the supernatants were quantified as a function of total insulin content from each sample.

### 2.10 Statistical analysis

All statistical analyses were performed using GraphPad Prism version 9.3.1 with a minimum of n = 3 biological replicates per sample group. Statistical comparisons were performed with two-tailed unpaired T-tests, or two-way ANOVAs where indicated, and were considered significant at p < 0.05.

## 3. Results

### 3.1 Sub-lethal DNA damage with etoposide leads to senescent phenotypes in NIT1 cells

To investigate whether DNA damage leads to senescence phenotypes in β cells relevant to T1D, we used the NIT1 β cell line, derived from an insulinoma in the T1D-susceptible NOD/LtJ mouse strain [21]. For comparison, we also examined the MIN6 β cell line derived from the non-T1D susceptible C57BL6 strain [20], which was recently shown to develop some senescent phenotypes in response to DNA damage with bleomycin [34], although a comprehensive investigation in this line was not performed. We induced DNA double-strand breaks using the topoisomerase II (TopoII) poison etoposide [35], as this drug has been used in the clinic to treat insulinomas [36] and leads to an increased burden of unrepaired endogenous DNA double-strand breaks due to inhibited ligation after cleavage by endogenous TopoII enzymes. Of the two type II topoisomerase family members, we found that TopoIIα was expressed in both MIN6 cells and NIT1 cells, while TopoIIβ was expressed only in MIN6 (**Supplementary Figure 1A**) confirming that these cells express the target of this drug.

MIN6 or NIT1 cells were cultured in etoposide for 3 days, followed by drug removal to assay early senescent phenotypes or culturing the cells in drug-free media for an additional 5-14 days to assay late senescent phenotypes (Figure 1A). We titrated the doses of etoposide to maintain viability throughout the treatment and culture period. While MIN6 cells tolerated 2 µM with >60% viability for up to 14 days post-drug removal, NIT1 could only survive in up to 0.5 µM for 7-9 days post-drug removal (Figure 1B) and similar results were obtained using 0.25 µM etoposide. Viability for both cell lines was maintained at ∼80-90% during 72 h of drug treatment (Figure 1B) but cell counts were significantly lower in the etoposide-treated cells as compared with controls (Supplementary Figure 1B), suggesting that while some cell death was occurring, there was also a proliferation arrest during drug treatment period.

**Figure 1.**
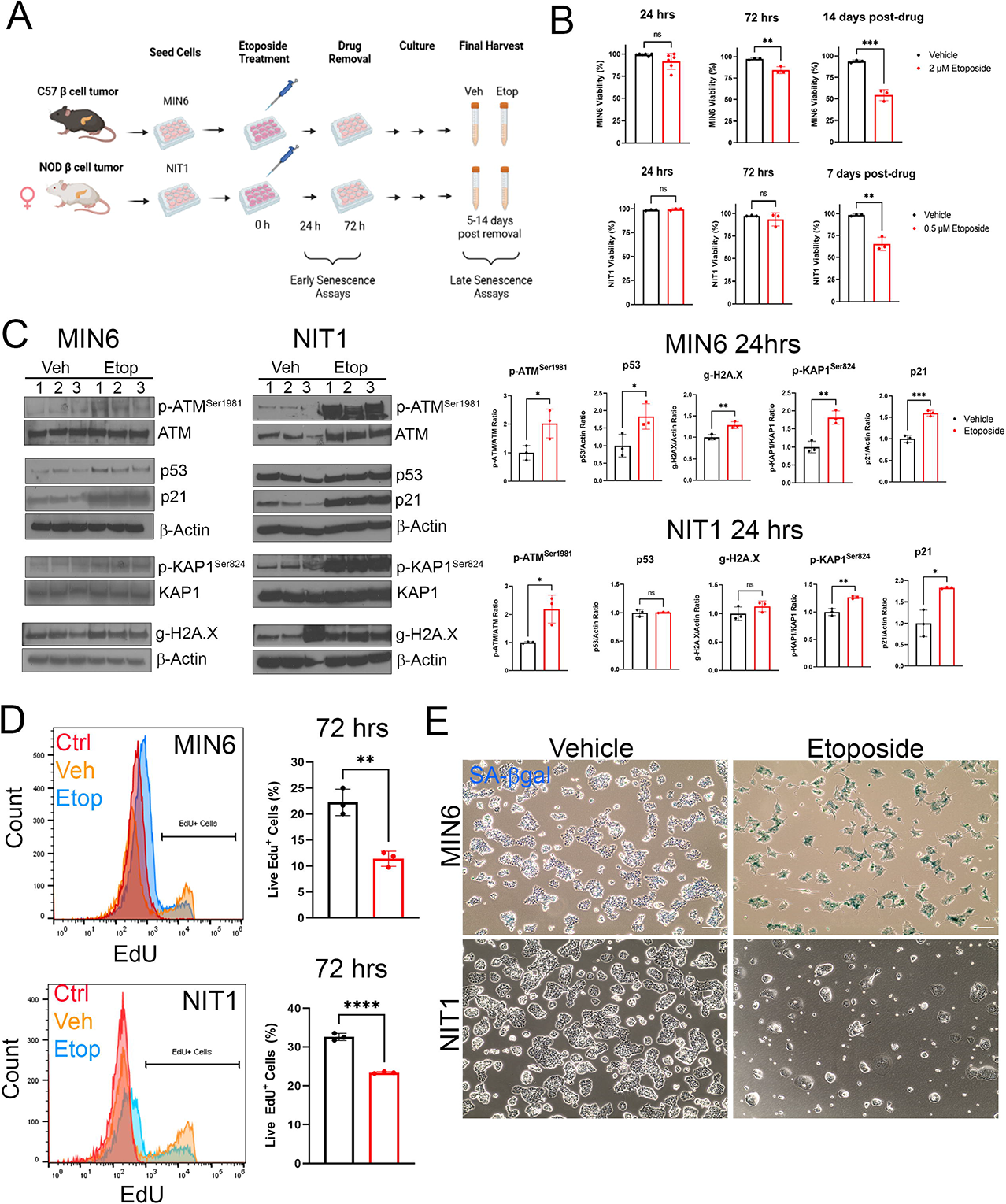
**Sub-lethal DNA damage with etoposide induces DDR, growth arrest and senescent morphology in MIN6 and NIT1 cells.** (A) MIN6 cells (derived from C57BL6 mouse beta cell tumor) or NIT1 cells (derived from NOD/LtJ mouse beta cell tumor) were seeded and treated with etoposide (2 µM for MIN6, 0.25-0.5 µM for NIT1) or vehicle control (DMSO) for 72 hrs. Drug was removed by changing cells into fresh media, and cells were cultured an additional 5-14 days before final harvests. Early senescence phenotypes were monitored at 24-72 hrs post-etoposide treatment and late phenotypes were assayed at 5-14 days post drug removal. B) Cell viability was scored at 24 or 72 hrs after etoposide treatment and 7 or 14 days post-drug removal. Data are mean ± SD of n = 3-6 biological replicates. C) Western blot analysis and relative quantifications of DDR activation markers in MIN6 and NIT1 cells at 24 h post etoposide or vehicle treatment. Data are mean ± SD of n = 3 biological replicates. D) EdU labeling flow cytometry assay for DNA replication at 72 hrs post-etoposide or vehicle control treatment of MIN6 or NIT1 cells. ‘Ctrl’ indicates a no EdU negative control population of cells. Data are mean ± SD of n = 3 biological replicates. E) Representative images of Xgal staining for SA-βgal activity in MIN6 and NIT1 cells at day 5 post-drug removal. Scale bar = 30 µm. *p < 0.05, **p< 0.005, ***p<0.0005, ****p < 0.00005, unpaired two-tailed T-tests.

β cells with DNA damage in vivo activate a canonical DDR involving the ATM-p53-21 pathway and phosphorylated KAP1^Ser824^ and phosphorylated Histone H2A.X^Ser139^ (gamma-H2A.X) [7,19,37,38]. Therefore, we next characterized DDR activation in MIN6 and NIT1 cells 24 h after etoposide treatment (Figure 1C). MIN6 cells exhibited a coordinated DDR with significant increases in phosphorylated ATM^Ser1981^, total p53, gamma-H2A.X, phosphorylated KAP1^Ser824^ and p21 (Figure 1C). In contrast, while NIT1 cells had increased activated ATM, phosphorylated KAP1 and induction of p21, there was no increase in total p53 or gamma-H2A.X (Figure 1C). Activated p53 (phospho-p53^Ser15^) was also detected in both undamaged and etoposide-treated MIN6 and NIT1 cells, but was not significantly increased relative to total p53 levels (Supplementary Figure 2). Total p53 and gamma-H2A.X were already abundant in undamaged NIT1 and MIN6 cells (Figure 1C). Etoposide treatment also reduced the percent of MIN6 and NIT1 cells replicating DNA as indicated by a lower proportion of cells labeled with EdU at 72 h post-drug treatment (Figure 1D) consistent with a growth arrest. Similarly, etoposide treated MIN6 cells developed an enlarged flattened morphology with strong SA-βgal activity by 5 days after drug removal, however, while NIT1 cells showed enlarged flattened morphology, they did not stain positive for SA-βgal activity at the same time point (Figure 1E). Taken together, these data show that NIT1 cells are more sensitive to DNA damage than MIN6 and that sub-lethal DNA double-strand break damage triggers canonical DDR activation, growth arrest and senescence-related morphology changes.

### 3.2 Cdkn1a but not Cdkn2a is upregulated in growth arrested NIT1 cells

We next assessed whether CDK inhibitors p21 and/or p16^Ink4a^ were upregulated at later stages after drug removal in NIT1 cells (Figure 2). Notably, at the 72 h post-treatment timepoint, *Cdkn1a* but not *Cdkn2a* was upregulated in MIN6 cells and this persisted into late stage senescence at 14 days post-drug removal (Figure 2A). Similarly, NIT1 cells also upregulated *Cdkn1a* expression at 72 h post-treatment and 7 days post-drug removal and also showed no upregulation of *Cdkn2a* (Figure 2B). Elevated p21 expression was also reflected at the protein level 72 h post-treatment and 7 or 14 days post-drug removal in both lines, whereas again p16^Ink4a^ was not detected (Figure 2C and 2D). We also assessed the expression of Bcl-2, which is selectively upregulated during β cell senescence in T1D and confers a prosurvival phenotype [7]. Notably, Bcl-2 was below the detection limit in control or etoposide treated MIN6 cells or NIT1 cells, despite robust expression in mouse testis as a positive control (Supplementary Figure 3). Taken together these data suggest that senescent growth arrest in NIT1 involves p21 but not p16^Ink4a^ and does not involve Bcl-2 upregulation.

**Figure 2.**
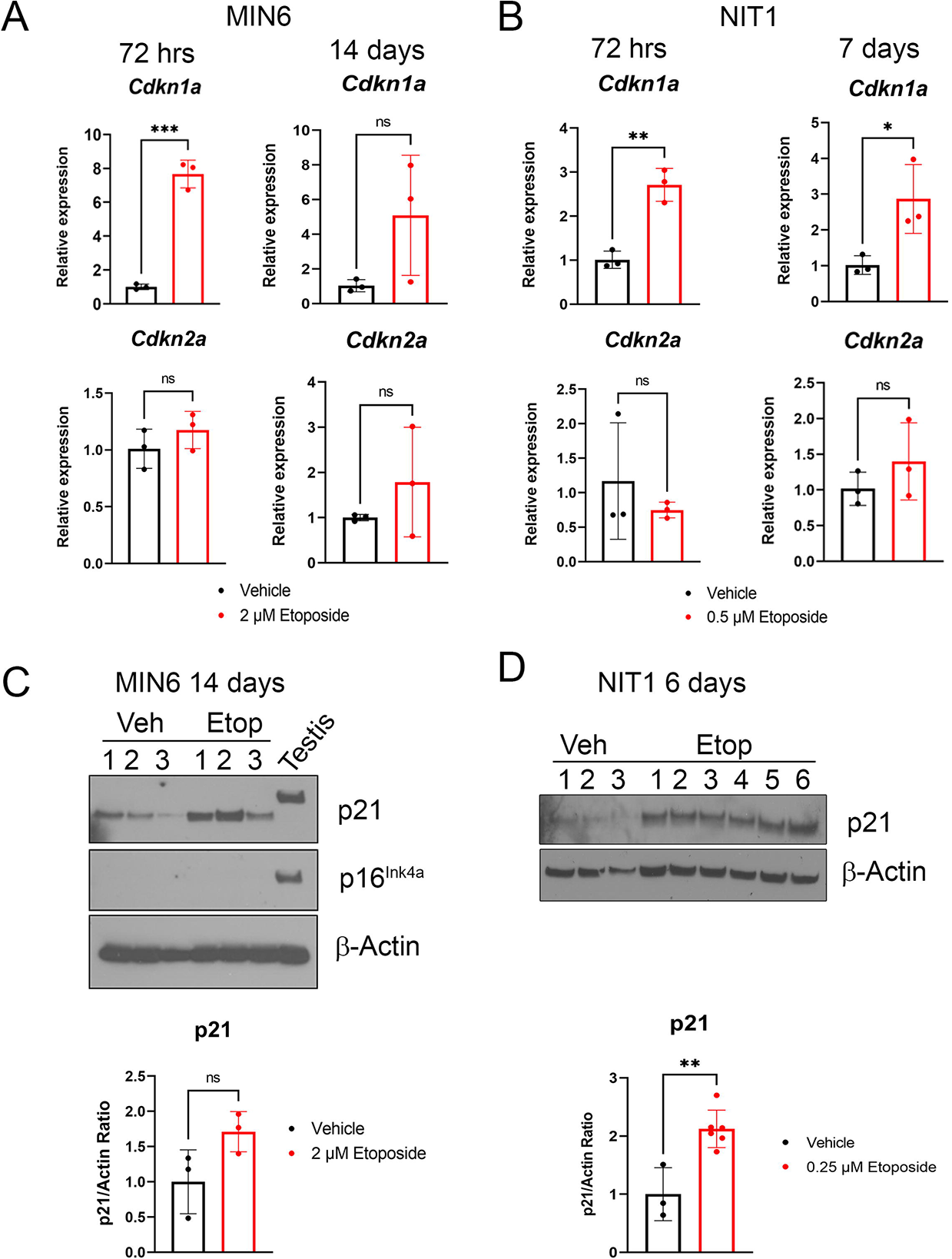
**Induction of *Cdkn1a* but not *Cdkn2a* at mRNA and protein level in etoposide treated MIN6 and NIT1 cells** A) qRT-PCR analysis of *Cdkn1a* and *Cdkn2a* expression normalized to *Gapdh* in MIN6 cells at 72 hrs post-etoposide or vehicle treatment or 14 days post-drug removal (as from Fig. 1). Data are mean ± SD of n = 3 biological replicates. B) qRT-PCR analysis as in B) except on NIT1 cells at 72 hrs and 7 days post-drug removal. C) Western blot of p21 and p16^Ink4a^ on whole cell extracts from MIN6 cells treated with 2 µM etoposide or vehicle control 14 days post-drug removal, Actin was a loading control and whole cell extract from 3 month old C57BL6 mouse testes was a positive control for p21 and p16. P21 quantification relative to actin is mean ± SD of n = 3 biological replicates. Data are mean ± SD of n = 3 biological replicates. D) Western blot of p21 on whole cell extracts from NIT1 cells treated with 0.25 µM etoposide or vehicle at 6 days post-drug removal. Data are mean ± SD of n = 3 biological replicates. For all panels, *p < 0.05, **p< 0.005, ***p < 0.0005, ns = not significant, two-tailed T-tests.

### 3.3 Differences in SASP development and UPR in MIN6 and NIT1 cells following DNA damage

SASPs are cell-type and stressor-dependent, dynamic programs of secreted cytokines, chemokines, matrix proteases, shed receptors, growth factors, microRNAs and extracellular vesicles that are immunogenic and exert the paracrine effects of senescent cells within the tissue microenvironment [39–41]. In senescent β cells during pathogenesis of T1D in NOD mice, SASP comprises a wide variety factors such as interleukins (IL-6), chemokines (Cxcl10), growth factors (Igfbp3), and matrix metalloproteases and protease inhibitors (Mmp2, Serpine1) [7, 14]. To determine whether etoposide-induced DDR in NIT1 cells led to a SASP resembling primary β cells in T1D, we assayed a panel of SASP factors we previously identified NOD mouse β cells by a combination of western blot and luminex secretion assays (Figure 3). Western blot of proenzyme and cleaved active Mmp2 isoforms revealed that etoposide treatment led to increased pro-Mmp2 levels in both MIN6 and NIT1 cells (Figure 3A and 3B). Luminex assays for SASP factors IL-6, Serpine1 and Igfbp3 [7] on conditioned media showed dramatically increased secretion of Serpine1 and Igfbp3, but not IL-6 from senescent NIT1 cells as compared with controls whereas none of these factors were secreted by late-stage senescent MIN6 cells (Figure 3C and 3D). Next, we determined whether induction of DDR and senescence would impact the UPR, a different β stress response implicated in T1D [4]. We validated antibodies for UPR mediators phospho-IRE1α ^Ser724^ (activated IRE1α) and total IRE1α, total PERK and total ATF6α (90 kDa uncleaved form) by western blot on primary human islets treated with 1µM Thapsigargin for 5 h, which led to increased levels of these UPR factors (Supplementary Fig. 4). Western blot analysis revealed induced p21 at 72 h post-etoposide treatment in both lines as expected, however, MIN6 cells also showed reduced phosphorylated and total IRE1α levels, whereas NIT1 cells showed no significant differences in any of these UPR mediators (Figure 3E and 3F), suggesting that DDR and early senescence leads to different effects on UPR mediators in MIN6 versus NIT1 cells. Taken together these data reveal that senescence induction leads to differences in SASP and UPR mediators between MIN6 and NIT1 cells.

**Figure 3.**
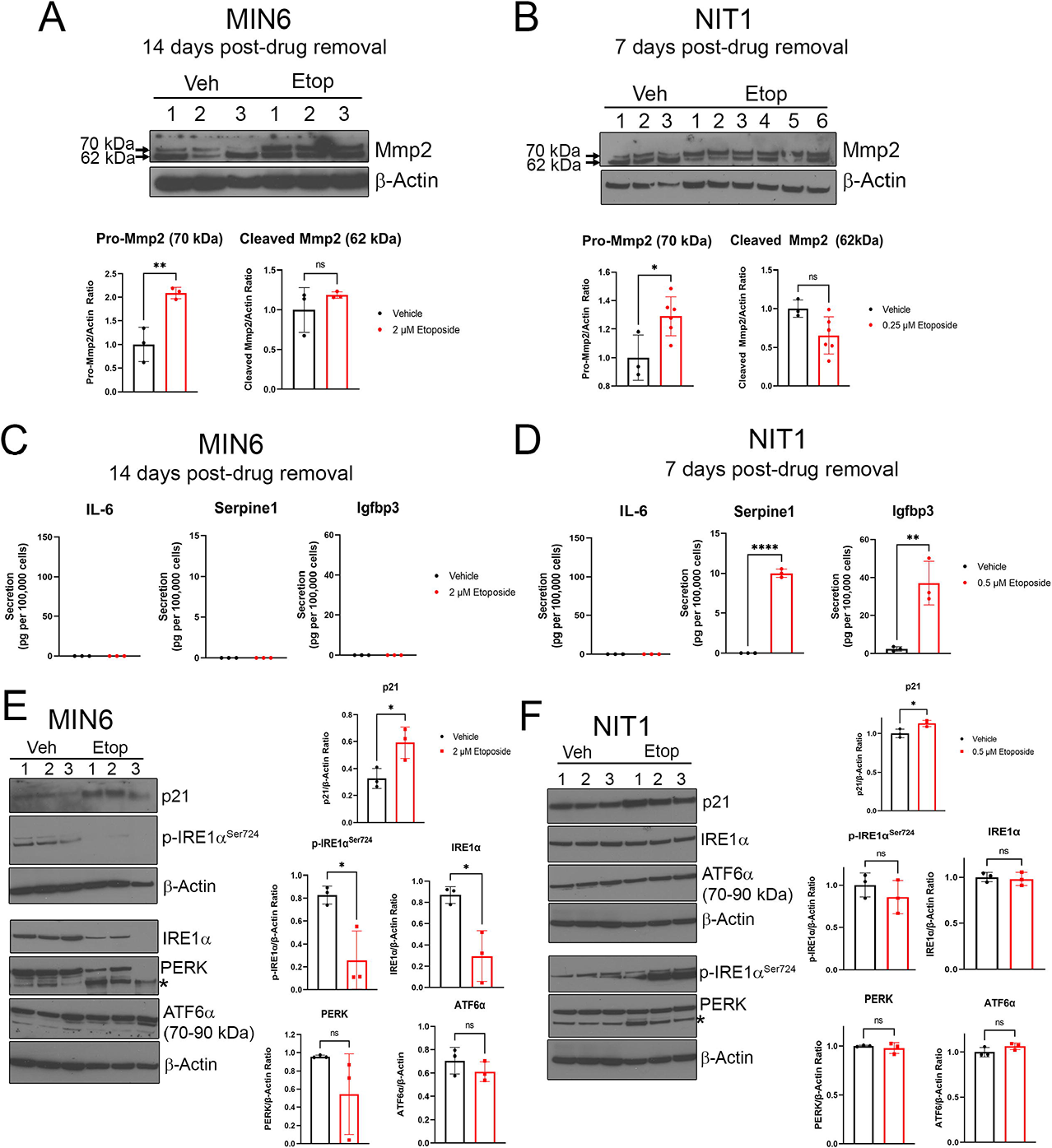
**Differences in SASP development and UPR in MIN6 and NIT1 cells following etoposide treatment.** A) and B) Western blot analysis of Pro-Mmp2 (70 kDa) and cleaved/activated Mmp2 (62 kDa) on whole cell extracts from MIN6 or NIT1 cells treated with vehicle or etoposide as indicated. Quantification shows relative Pro-Mmp2 or Cleaved Mmp2 to Actin, mean ± SD of n = 3 biological replicates. C) and D) Luminex assay of secreted SASP factors IL-6, Serpine1 and Igfbp3 in serum-free conditioned media from MIN6 or NIT1 cells treated as indicated at timepoints indicated. Data are mean ± SD from n = 3 biological replicates. E) and F) Western blot analysis of UPR mediators phosphorylated and total IRE1α, total PERK and total ATF6α in MIN6 and NIT1 cells at 72h post-treatment with etoposide. p21 levels were used as a positive control and β-Actin was a loading control. Data are mean ± SD of n = 3 biological replicates. Asterisk on the blots indicates a non-specific band for the PERK blots. For all panels, *p< 0.05, **p<0.005, ***p<0.0005, ns = not significant, two-tailed T-tests

### 3.4 RNA-seq reveals DDR and senescence transcriptional program of human islets

To provide an unbiased analysis of the transcriptional program following DDR and senescence in human islets, we next performed RNA-seq on senescent and control human islets cultured ex vivo using our previous human islet senescence model [7, 14] (Figure 4). Primary islets isolated from a healthy female donor were cultured with bleomycin to induce DNA damage for 48 h and total islet RNA from the samples was extracted at 4 days post-drug removal for paired-end RNA-seq (Figure 4A). This timepoint is typically 1 day prior to the detection of SASP secretion in this model [7, 14], and we reasoned that transcriptional changes in SASP genes would precede increased protein secretion, as SASP is controlled transcriptionally [14, 15]. Differential expression analysis revealed 1690 genes significantly downregulated ≥0.5-fold and 201 genes significantly upregulated ≥2-fold (p < 0.05) (Figure 4B). A complete list of all genes with detected expression levels as fragments per kilobase per million reads (FPKM) are listed in Supplementary Table 1. Differentially expressed genes were enriched for gene ontology (GO) terms including ‘cell division’ and ‘cell cycle’ and KEGG pathways ‘cell cycle’, ‘p53 signaling pathway’ and ‘PI3K-Akt signaling pathway’ (Figure 4C, Supplementary Figure 5A). Genes involved in proliferation and cell cycle progression such as *MIK67, RRM2, TOP2A, CCNA2*, *CCNB2, MCM10, CDK1* were significantly downregulated (Figure 4D) and p53 DDR pathway genes including *SPATA18, DDB2, GADD45A, GADD45G, ACER2, ZMAT3* [42] were significantly upregulated (Figure 4E). In addition, p53 target gene *CDKN1A* was upregulated, but not other cyclin-dependent kinase inhibitor genes such as *CDKN2A*, *CDKN1B*, *CDKN1C, CDKN1D* or *CDKN2B* (Figure 4E **and Supplementary Table 1**). Gene expression changes associated with early senescence also occurred, including downregulation of *LMNB1* and *HMGB2* (Figure 4E) both of which occur during p53-mediated senescence induction in other cell types [43, 44]. In addition to mRNAs, we also noted several small noncoding RNAs that were differentially expressed. These included *MIR34A*, a microRNA involved in senescence [45], along with *MIR3687-2*, *MIR3648-2, SNORA46 SNORA84, SNORD94* and *SNORD14B* which were all upregulated and *MIR22, MIR222, MIR143, MIR1-2*, *DNMOS3* were downregulated (**Supplementary Table 1**). We also found a group of previously annotated long noncoding RNAs (lncRNAs) that were differentially expressed and a group of predicted novel lncRNAs in bleomycin-treated islets (**Supplementary Table 2**). Notably among the lncRNAs, *DINOL* was found to be upregulated (**Supplementary Table 1**), consistent with its established role in amplifying p53-p21-mediated DDR signaling [46]. Together, these data validate that bleomycin treatment activates a p53-p21 DDR and early senescence transcriptional program in adult human islets.

**Figure 4.**
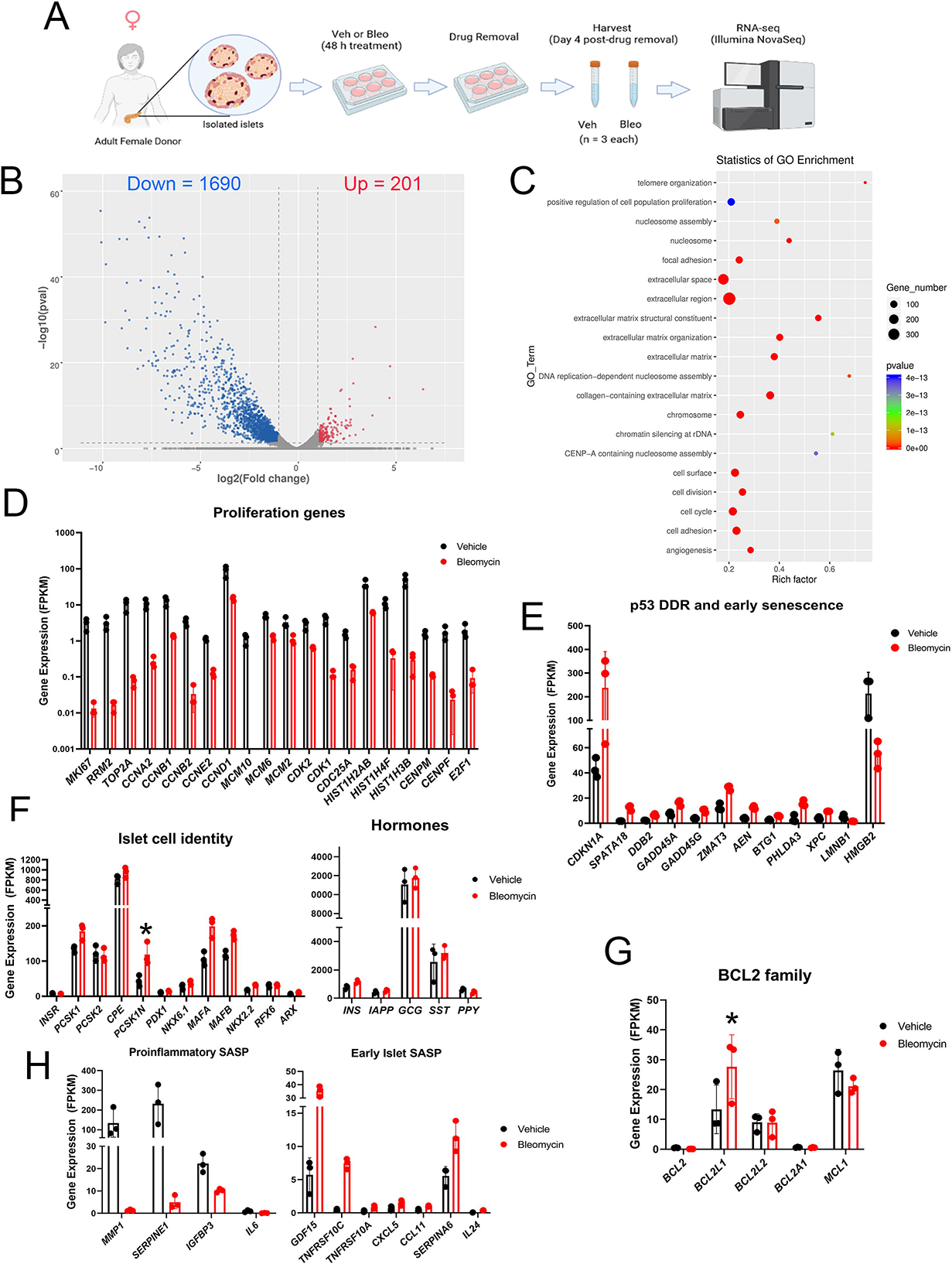
**RNA-seq analysis reveals activation of the p53-p21 transcriptional program during DDR-mediated senescence in human islets.** A) Overview of human islet DDR and senescence model. Islets isolated from a 44 year-old female donor were rested overnight and then divided into a total of 6 wells and cultured in the presence of vehicle (DMSO) or 50 µM bleomycin for 48 h (n = 3 biological replicates per group). The islets were then transferred to fresh drug-free media and cultured an additional 4 days prior to harvesting on day 5 post-drug removal for RNA extraction and paired-end RNA-seq. B) Volcano plot of differentially expressed genes (p < 0.05) indicating 1690 genes downregulated ≤0.5-fold and 201 genes upregulated ≥2-fold in bleomycin treated islets C) Significant GO terms of differentially expressed genes. D) Plot of normalized expression levels (FPKM) of selected significantly downregulated proliferation and cell cycle genes. E) Plot of normalized expression levels of significantly upregulated p53-p21 DDR genes and two early senescence genes (*LMNB1* and *HMGB2*). F) Plot of normalized expression levels of selected islet cell identity and hormone genes. None of these genes were differentially expressed except *PCSK1N* which was significantly upregulated in bleo versus control (*p < 0.05). G) Plot of selected BCL-2 family anti-apoptotic genes. Only *BCL2L1* was significantly upregulated, (*p < 0.05). H) Plot of selected significantly downregulated proinflammatory SASP genes and significantly upregulated SASP genes.

### 3.5 DDR and senescence does not alter human islet cell identity genes or UPR genes

To assess the effects of DDR activation and senescence on human β cell/endocrine islet cell function and identity we surveyed genes involved in hormone production and endocrine cell identity (Figure 4F). Genes encoding endocrine islet cell hormones and processing enzymes (*INS, GCG, SST, INSR, PCSK1, PCSK2, CPE*) and key beta/islet cell transcription factors (*PDX1, NKX6.1, MAFA, MAFB, NKX2.2, RFX6, ARX*) were not changed (Figure 4F). The only exception was upregulation of *PCSK1N*, which encodes proSAAS, an endogenous inhibitor of the PCSK1 hormone processing enzyme that mediates proinsulin cleavage [47] (Figure 4F). Furthermore, genes involved in ER stress and UPR (*XBP1, DDIT3, EDEM1, ERN1, ATF4, ATF6, HSPA5, HSP90B1*) [4], were not significantly changed between control and bleomycin treatment (Supplementary Figure 5B). Comparison of the list of differentially expressed genes in our dataset with recently generated human islet RNA-seq datasets during aging [48] and adult human islets treated with the cytokine IFNα [49] showed that there were only 74/385 (∼19%) age-related genes in common and only 256/1894 (∼13%) IFNα-regulated genes in common, respectively, (Supplementary Figure 5C, Supplementary Table 1). This suggested that the transcriptional response during islet DDR activation and senescence is generally different from natural aging and cytokine exposure.

### 3.5 BCL-2 family genes and SASP genes in islet DDR and senescence

To assess whether islet cells were developing late senescence phenotypes, we next looked at BCL-2 family prosurvival genes and SASP genes. Among the prosurvival family members, *BCL2L1, MCL1* and *BCL2L2* were the only members expressed at appreciable levels in islets, and of these only *BCL2L1* (encoding BCL-XL) was significantly upregulated (Figure 4G). Assessment of genes encoding canonical proinflammatory SASP factors such as *IL6, CXCL8, SERPINE1, IGFBP3, CXCL1, IGFBP4, MMP1, MMP2* revealed that the majority of them were either dramatically downregulated or unchanged in bleomycin treated islets relative to controls (**Supplementary Table 1**). However, among the significantly upregulated genes, we noted several previously identified SASP genes encoding secreted factors and cell surface receptors, including *GDF15, CXCL5, CCL11, TNFRSF10C, TNFRSF10A* and *SERPINA6* [16,34,39,50,51] (Figure 4H).

### 3.6 Bleomycin induces DDR and senescence phenotypes in a human β cell line model

To corroborate the p53-p21 pathway transcriptional changes identified by RNA-seq on islets and determine whether increased SASP gene expression is reflected in protein secretion, we carried out western blot for DDR and senescence proteins and Luminex assays on two independent islet preparations at day 5 post-drug removal isolated from adult male donors (Figure 5A). Notably, westerns confirmed significantly increased expression of DDR markers phospho-ATM^Ser1981^ and p21 (Figure 5B). In addition, Luminex assays revealed increased secretion of previously characterized islet SASP factors IL-8, CXCL1 and IGFPB4 along with newly identified factors TNFRSF10C and GDF-15 (Figure 5C). Since islets are a mixed cell population, the direct contribution of β cells to the observed senescent phenotypes cannot be determined. Therefore, we turned to the EndoC-βH5 human β cell line model to determine whether similar phenotypes could be induced in directly in human β cells experiencing DNA damage (Figure 5D). EndoC-βH5 cells are the latest derivative of the widely used EndoC human fetal β cell lines [23] that has been clonally selected for maximal Insulin secretion and β cell identity markers and is essentially non-proliferating. After seeding and 2 days in culture, we treated EndoC-βH5 cells for 48 h with vehicle (0.1% DMSO) or 35 µM bleomycin, followed by drug removal and additional culture for 2-5 days to monitor early and late senescence phenotypes (Figure 5D). Notably, 48 h after bleo treatment, these cells activated DDR signaling as judged by western blot analysis for phospho-ATM^Ser1981^, gamma-H2A.X and p21 (Figure 5E). QRT-PCR demonstrated a sustained expression of *CDKN1A* 2 days after drug removal, but again *CDKN2A* was unaffected (Figure 5F). We also confirmed upregulation of SASP and BCL-2 family genes (*CXCL8* and *BCL2L1*) (Figure 5F) suggesting development of late senescence phenotypes starting at 2 days post-drug removal. To determine whether SASP factor secretion was occurring we monitored secretion of islet SASP factors IL-8, IGFPB4, CXCL1 and newly identified factors TNFRSF10C and GDF-15 by Luminex on conditioned media at day 5 post-drug removal. Of these factors, TNFRSF10C was not detectable and only GDF-15 showed significantly increased secretion (Figure 5G). Taken together, these data suggest that DNA damage is sufficient to induce senescent phenotypes directly in human β cells.

**Figure 5.**
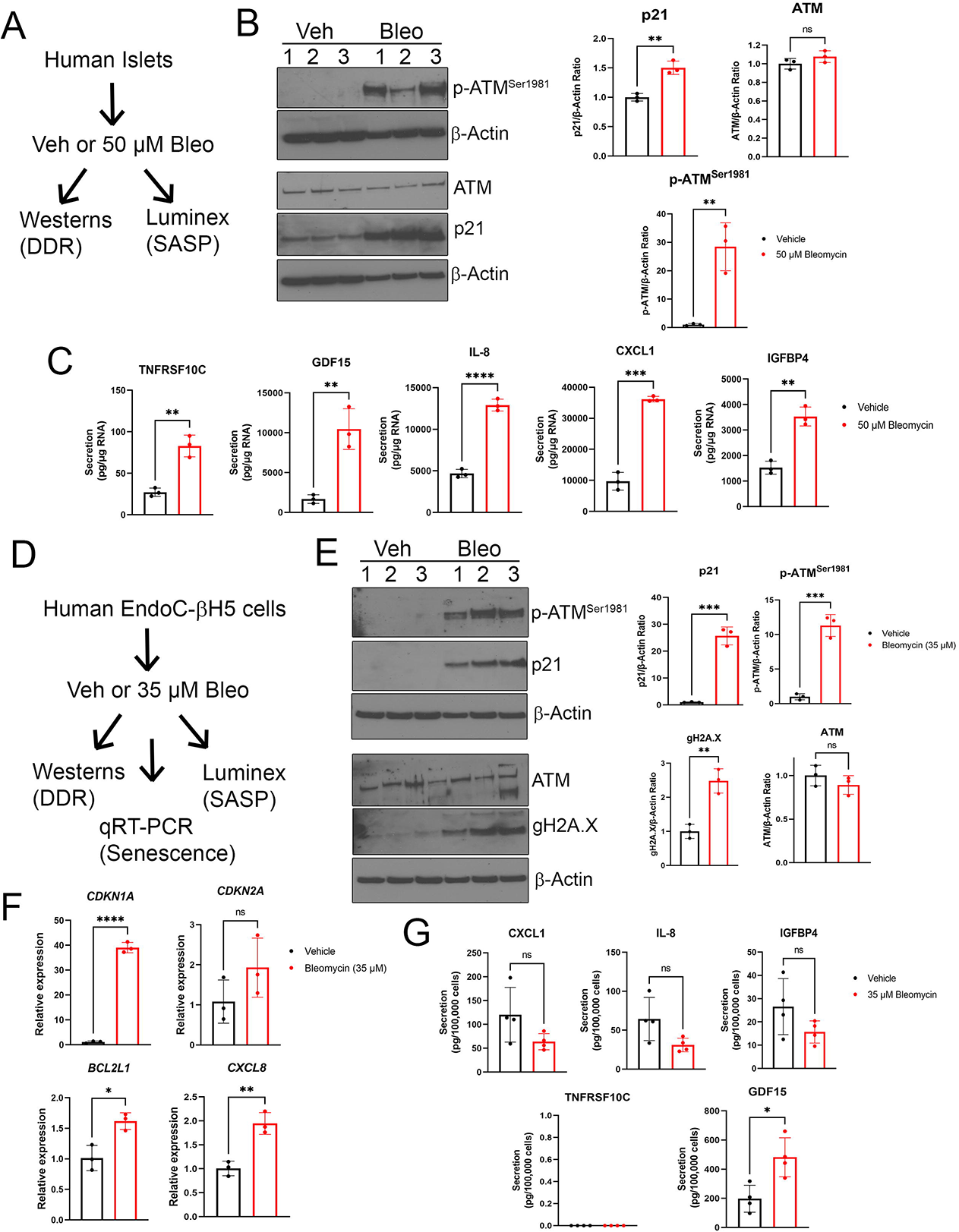
**Validation of DDR and SASP activation in human islets and the EndoC-βH5 human β cell line.** A) Overview of experiment on human islets. Islets from adult male donors (ages 32 or 55) were treated as indicated and harvested for western blot and Luminex assays. B) Western blot analysis of DDR and senescence proteins on vehicle (DMSO) and bleomycin-treated islets at day 4 post-drug removal. Data are means ± SD of n = 3 biological replicates. C) Luminex assays for indicated SASP factors in the conditioned media from islets treated as in (A) at day 5 post-drug removal. Data are normalized to islet RNA content and are means ± SD of n = 3 biological replicates. D) Overview of experiment on EndoC-βH5 cells. Cells were treated as indicated and harvested for western blotting, qRT-PCR and Luminex assays. E) Western blot analysis of DDR and early senescence markers in cells treated as indicated. Data are means ± SD of n = 3 biological replicates. F) qRT-PCR analysis of senescence genes *CDKN1A, CDKN2A*, prosurvival gene *BCL2L1* (encoding BCL-XL) and SASP gene *CXCL8* (encoding IL-8) in EndoC cells treated as in (A) at day 2 post-drug removal. Data are means ± SD of n = 3 biological replicates. G) Luminex assays of indicated SASP factors in the conditioned media from EndoC cells treated as in (A) at day 5 post-drug removal. Data are normalized to viable cell counts and are means ± SD of n = 4 biological replicates. For all panels, *p<0.05, **p<0.005, ***p<0.0005, two-tailed T-tests.

### 3.7 Senescence decreases β cell insulin content

Finally, we explored the effect of DDR and senescence on insulin secretion (Figure 6). Surprisingly, induction of DDR and senescence in MIN6 and NIT1 cells led to abnormally increased GSIS with elevated basal and glucose-stimulated insulin secretion (GSIS) from both lines (Figure 6A). Early senescent NIT1 and MIN6 cells also showed significantly reduced insulin content (Figure 6A). In contrast, established senescence at day 5 post-drug removal in human islets did not affect GSIS, but led to decreased insulin content (Figure 6B). Taken together, these results suggest that DDR and senescence diminishes insulin content and shows different on GSIS in mouse β cell lines as compared with human islets.

**Figure 6.**
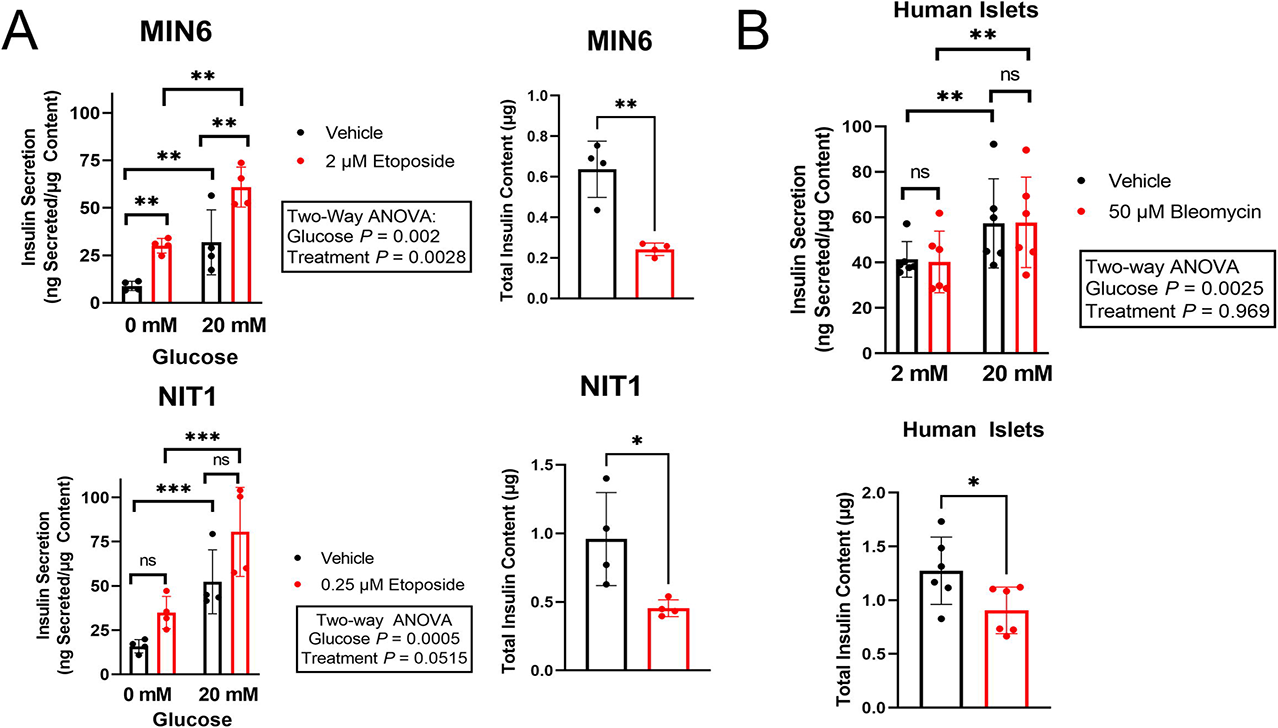
**Senescence leads to decreased insulin content.** A) GSIS assays and total insulin content of MIN6 and NIT1 cells treated with vehicle or etoposide at 72 h post-treatment. Data are means ± SD of n = 4 biological replicates. B) GSIS assays and total insulin content of human islets from a 55 year-old male donor treated with vehicle or bleomycin at day 4 post-drug removal. Data are means ± SD of n = 6 biological replicates. For GSIS panels, *p<0.05, **p<0.005, ***p<0.0005, Two-way ANOVAs. For Insulin content panels, *p< 0.05, **p<0.005, two-tailed T-tests.

## 4. Discussion

A subpopulation of β cells acquire senescence as a stress response during development and onset of T1D [7], but the triggers/stressors that lead to senescence in this setting are not known. The use of cell culture models has been indispensable for the field of senescence and has been foundational for understanding the mechanisms underpinning this complex process. Since senescent β cells in T1D show DDR activation [7, 19] and persistent unrepaired DNA damage is a known inducer of senescence, we carried out studies to determine the extent to which chemically induced DNA damage in the NIT1 cell line derived from the T1D-susceptible NOD mouse strain, adult human islets and EndoC cells recapitulates features of β cell senescence in T1D. We demonstrated that NIT1 cells responded to accumulated endogenous DNA double-strand break damage from low concentrations of etoposide by activating the DDR, growth arrest, developing an enlarged flattened morphology and a SASP resembling what is observed in T1D. There were some notable differences in these phenotypes as compared with MIN6 cells, which did not secrete most of these SASP factors, pointing to the role of genetic background in modifying senescent phenotypes. Consistent with previous findings revealing polymorphisms that reduce the efficiency of DNA double-strand break repair in NOD/ShiLtJ as compared with C57BL10 embryonic fibroblasts [18], NIT1 cells were more sensitive to DNA double-strand break damage as compared with MIN6. In addition, although they were clearly growth arrested and developed an enlarged and flattened morphology, NIT1 cells also did not develop high SA-βgal activity like MIN6 cells and β cell senescence during T1D [7]. Both cell lines had sustained upregulation of *Cdkn1a* expression at the gene and protein levels but not *Cdkn2a*, suggesting that the growth arrest relies on p21 rather than p16^Ink4a^. Furthermore, Bcl-2 was not detected in NIT1 or MIN6 cells and consistent with a lack of prosurvival phenotype, these lines showed declining viability in culture at late stages. Taken as a whole, these results suggest that specific aspects of mouse β cell senescence in T1D, including DDR activation, growth arrest and SASP, but not SA-βgal activity or Bcl-2 prosurvival phenotype can be modeled using NIT1 cells treated with sub-lethal concentrations of etoposide.

Human islets show substantial differences in architecture, metabolism, and proliferative capacity as compared with rodents [52], necessitating the study of β cell DDR and senescence in human models. We previously established a primary human pancreatic islet model for DDR and senescence using the DNA double-strand break agent bleomycin and found that it induces some of the same phenotypes observed during β cell senescence in human T1D donors [7]. However, a systematic interrogation of this model was lacking and it was previously not determined whether β cells contributed to the observed phenotypes. Using an unbiased analysis by RNA-seq, we found that DDR activation and senescence induction by bleomycin treatment of islets resulted in a coordinated p53-p21 transcriptional program, involving downregulation of genes involved in proliferation and cell cycle progression and upregulation of p53-targeted genes. Although there were a small subset of genes in common with other human islet RNA-seq datasets representing natural aging [48] or cytokine exposure [53] the changes during islet DDR activation and senescence were largely distinct. Importantly, the changes in this model occur 4 days after drug removal (6 days post-treatment) and thus reflect a stable phenotypic change rather than an acute response. Notable among these changes was the robust upregulation of *CDKN1A* (p21) and no increases in *CDKN2A* (p16) expression, consistent with our earlier analysis by qRT-PCR and protein-level changes in senescent β cells of T1D donors [7]. We further corroborated these changes in a human β cell line model, EndoC-βH5 cells, confirming that it can occur directly in β cells. While *CDKN2A* gene expression was not induced in islets after DDR and senescence, p16^INK4A^ protein level is already very abundant in middle-aged adult donor pancreas and islets [7,54,55], thus the gene expression level is likely not an accurate reflection of the protein levels. Remarkably, there were no significant changes in islet cell hormone genes, lineage identity genes, or UPR genes at this timepoint after DDR activation and early senescence. In addition to a p53-p21 transcriptional response in bleomycin-treated islets, we also found evidence for two other senescence phenotypes, BCL-2 family gene upregulation and SASP. We identified selective upregulation of BCL2 family member gene *BCL2L1* in bleomycin treated islets. The BCL-2 family of anti-apoptotic proteins confer a prosurvival phenotype on senescent cells and *BCL2L1* encodes BCL-XL, which is known to be upregulated at the mRNA and protein level in other cell types during senescence [56, 57]. Thus, it will be interesting to investigate whether BCL-XL is upregulated during β cell senescence in T1D donors and confers a prosurvival phenotype to senescent human β cells.

While many of the classical proinflammatory SASP genes were downregulated during DDR-mediated senescence in islets, a subset of previously identified TGFβ-related SASP genes encoding secreted factors and shed receptors [58] were significantly upregulated at the RNA level. Notable among them were *GDF15* and *TNFRSF10C*, which were also validated at the protein level in secretion assays to monitor SASP. GDF15 was recently identified as a β cell SASP factor in mice in a T2D model [34] and an anti-apoptotic factor in human islets, conferring protection from cytokine-induced apoptosis [59]. Importantly, GDF15 has also been linked to age-related senescence and was recently suggested as part of a biomarker panel for senescence-related inflammation [39]. Given its potential role as an anti-apoptotic factor, it will be of interest to determine whether GDF15 confers a prosurvival phenotype on senescent β cells and/or modulates immune-mediated cytoxicity of β cells. Additionally, we found TNFRSF10C, which is a putative decoy receptor to be upregulated both at the RNA and protein levels and secreted following DDR and senescence induction from islets. Given the role of this receptor in inhibiting extrinsic apoptosis in the context of cancer [60], its increased secretion from senescent islet cells may play a role in protecting these cells from immune attack. Thus, we demonstrated that transcriptional upregulation of islet SASP components such as TNFRSF10C and GDF15 is linked with increased SASP secretion. Interestingly, when our islet SASP panel (IL-8, CXCL1, IGFBP4, TNFRSF10C, GDF15) was interrogated in an EndoC human β cell line model for DDR and senescence, we only found GDF15 to be increased in secretion. This may be due to a variety of reasons, such as: 1) differences in β cell maturity between human islet β cells and EndoC cells, 2) effects of different genetic background polymorphisms and/or 3) secretion from non-β cells in the islet. Further investigations will be required to catalogue the similarities and differences between SASP in human islets as compared with EndoC β cell lines and it would be advantageous to sort purify β cells from human islet undergoing DDR and senescence to more closely quantify differences in secretion.

Using our mouse β cell lines and human islet senescence culture models, we determined whether insulin secretion was altered by DDR activation and senescence. Previous work has shown that age-related senescence improves β cell function [54]. However, treatment of euglycemic NOD mice with senolytic compounds to remove senescent β cells does not significantly affect glucose tolerance in vivo or static GSIS on isolated islets ex vivo [7]. In contrast, treatment of T2D mouse models with senolytic compounds improves β cell function [13], although there is likely to be additional effects on senescent cells in other tissues in T2D models [61]. These studies suggest a variety of potential effects of senescence on β cell insulin secretory function, depending on the context and type of senescence. In the present study, GSIS assays revealed an abnormal hypersecretory response during DDR and early senescence in MIN6 and NIT1 cells, where both basal and stimulated insulin secretion was elevated, concomitant with diminished insulin content. Surprisingly, GSIS was not significantly impaired at the day 5 timepoint of DDR-mediated senescence in human islets. However, a decreased insulin content phenotype was apparent, similar to MIN6 and NIT1 cells. *INS* mRNA levels were not significantly different between control and senescent islets as determined by RNA-seq, suggesting that the decreased insulin content may be due to post-transcriptional changes, such as crinophagy or selective insulin granule degradation [62]. Nevertheless, the diminished insulin content phenotype during senescence may be reflective of a progressive functional impairment. Further studies will be required to interrogate the mechanisms involved in these β cell functional changes during DDR and senescence. In conclusion, our findings generally substantiate sub-lethal DNA double-strand break damage in mouse NIT1 cells and human islets and EndoC-βH5 cells as culture models for some of the key phenotypes of β cell senescence during T1D. Despite their limitations, these models will be useful for further dissecting the molecular mechanisms and functional consequences of senescence and its contributions to the pathogenesis of T1D.

## Acknowledgements.

The authors thank Dr. Anil Bhushan at University of California San Francisco (UCSF) supported by NIH R01DK121794 and R01DK118099, for providing IIDP human donor islets and resources for PJT to carry out the RNA-seq on human islets while a postdoctoral fellow.

## CRediT authorship contribution statement

**Gabriel Brawerman:** Conceptualization, Methodology, Investigation, Writing - review & editing. **Jasmine Pipella:** Methodology, Investigation, **Peter J. Thompson:** Conceptualization, Formal analysis, Methodology, Investigation, Resources, Writing - original draft, Writing - review & editing, Visualization, Supervision, Funding acquisition.

## Declaration of Competing Interest

The authors declare that there are no conflicts of interest.

## Availability of Data and Materials

The RNA-seq raw data and processed data files described in this study are publicly available on NCBI GEO Accession number GSE176324. Expression data tables are also available in Supplementary Information.

## Funding

This study was supported by start-up funds awarded to PJT from the University of Manitoba and the Children’s Hospital Research Institute of Manitoba and a grant from the University of Manitoba Research Grants Program.

## Notes

https://www.ncbi.nlm.nih.gov/geo/query/acc.cgi?acc=GSE176324

### Competing Interest Statement

The authors have declared no competing interest.

### Summary of Updates

1. Figure 3 was modified to include new results. 2. New results are included in Figures 5 and 6.

