## Supplementary Figures for "DNA damage models phenotypes of β cell senescence in Type 1 Diabetes"

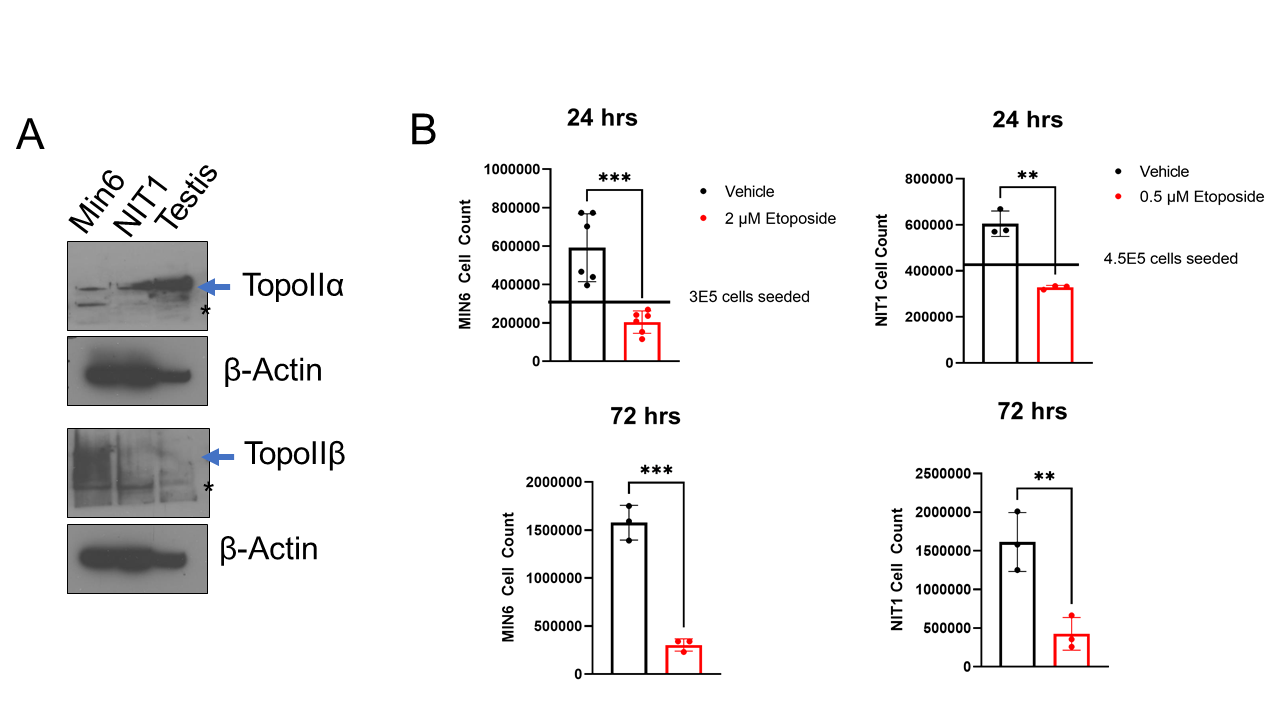

**Supplementary Figure 1. Etoposide treatment of MIN6 and NIT1 cells and effects on proliferation.** (A) Western blot analysis of Topoisomerase II family members in MIN6 and NIT1 whole cell extracts. Beta actin was a loading control and whole cell extract from 3 month C57BL6 testis was a positive control for TopoIIα. (B) Cell counts related to Figure 1B, showing viable cell numbers 24 or 72 h post-etoposide treatment. Cells were seeded at numbers shown and 24 h later treated with indicated concentrations of etoposide and then viable cells counted at timepoints indicated post-etoposide. Data are mean± SD of n = 3-6 biological replicates. **p <0.005, ***p<0.0005, two-tailed T-tests.

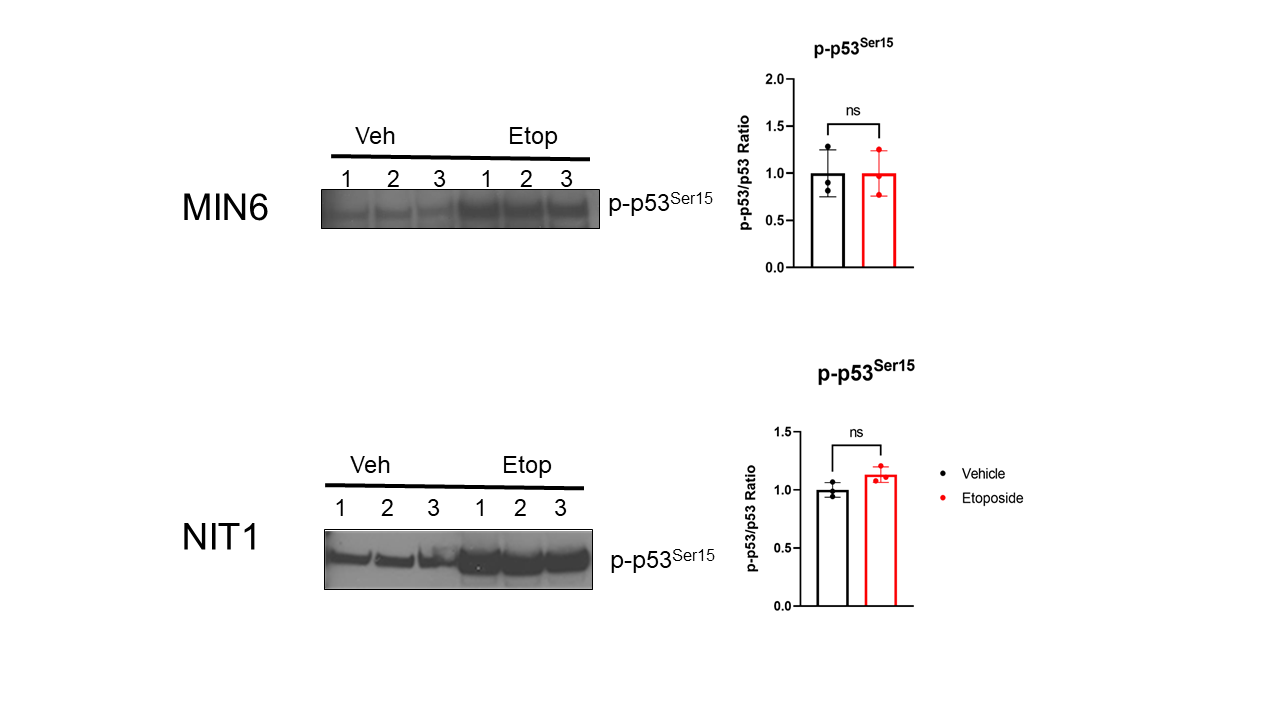

**Supplementary Figure 2. Western blot of phospho-p53^Ser15^ in MIN6 and NIT1 cells.** Westerns for p-p53^Ser15^ on control or Etoposide treated MIN6 and NIT1 cells 24 h after exposure, as from Figure 1C. p-p53 bands were normalized to p53 blots for each cell line shown in Figure 1C. Data are means ± SD from n = 3 biological replicates. Ns = not significant.

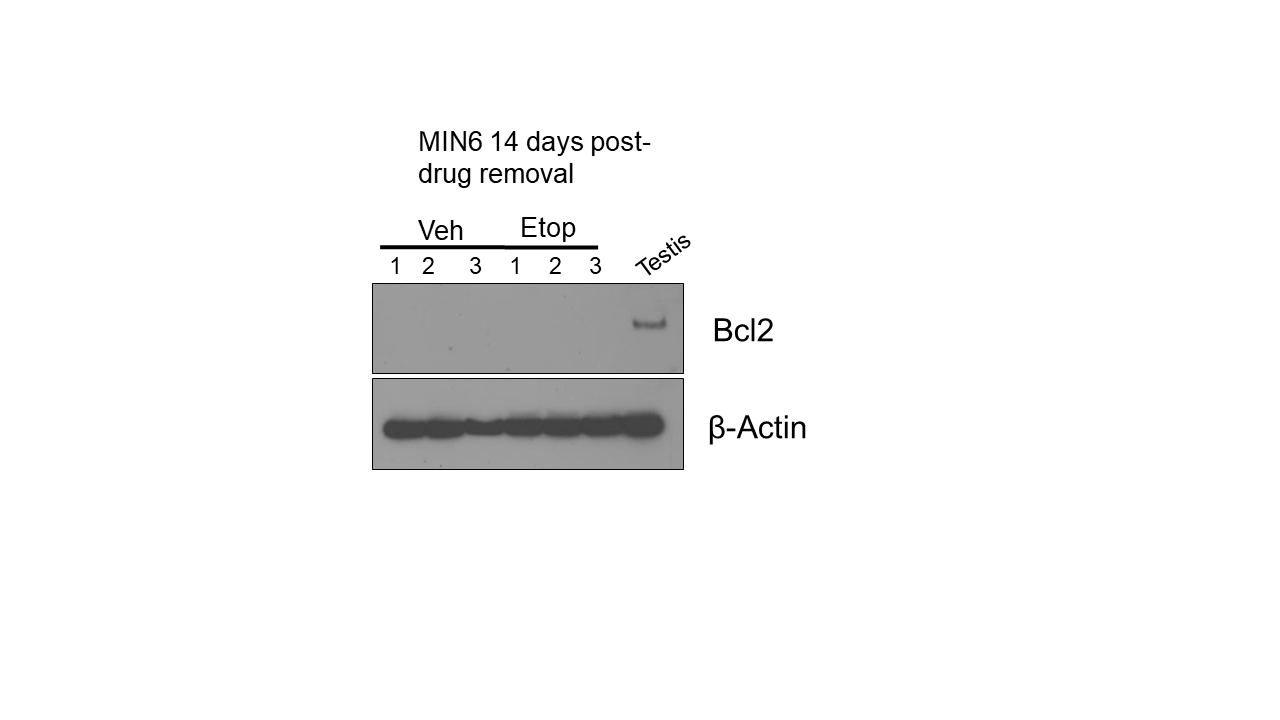

**Supplementary Figure 3. Bcl-2 protein levels in MIN6 cells.** Western blot was performed for Bcl-2 on whole cell extracts from MIN6 cells (vehicle control or 2 µM etoposide treated), at indicated timepoint, as from Figure 2C. Testis extract was a positive control and Beta actin was a loading control. Lack of Bcl-2 protein was also observed in control and etoposide treated (0.25 µM) NIT1 cells at day 6 post-drug removal (not shown).

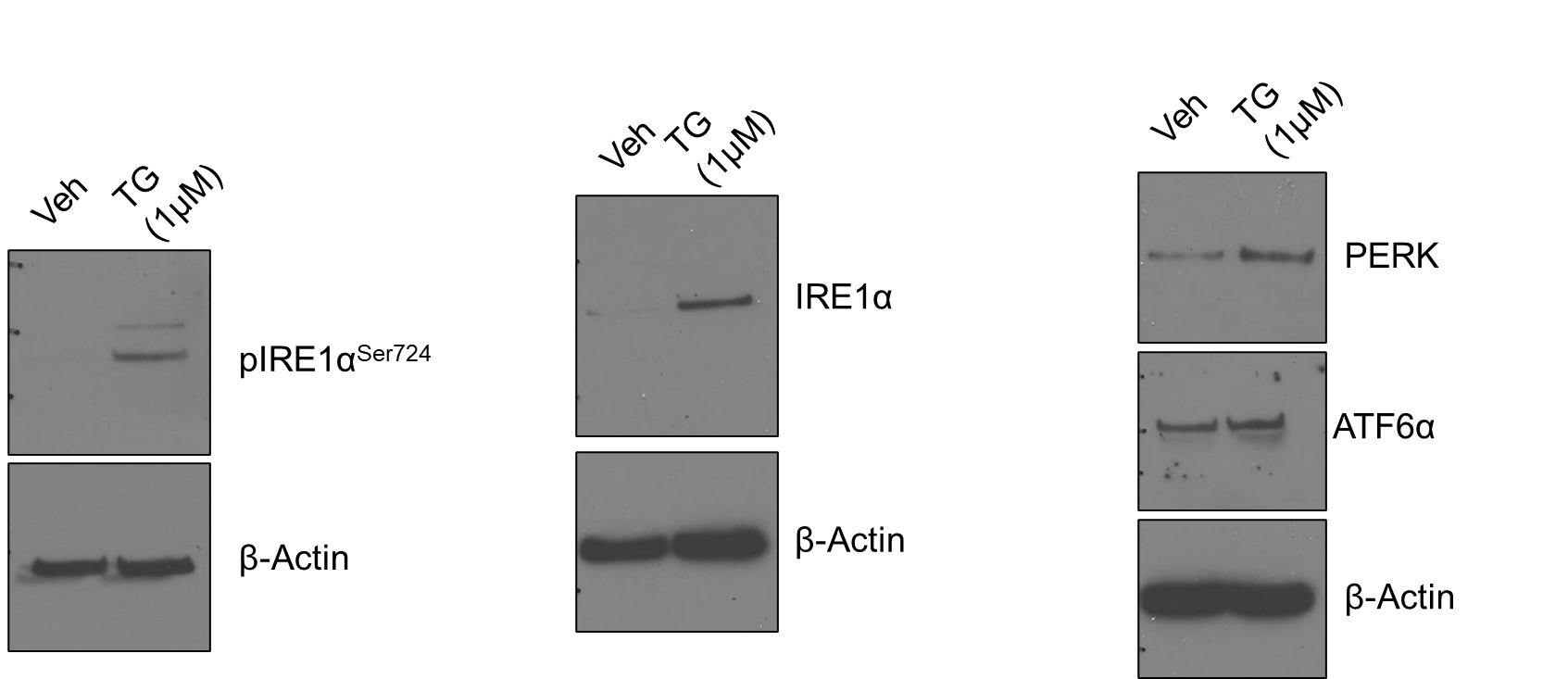

**Supplementary Figure 4. Validation of UPR antibodies on human islets.** Antibodies for phospho-IRE1α^Ser724^, total IRE1α, PERK and ATF6α on primary human donor islets treated with vehicle (0.1% DMSO) or 1μM Thapsigargin (TG) for 5 h. β-Actin was detected as a loading control.

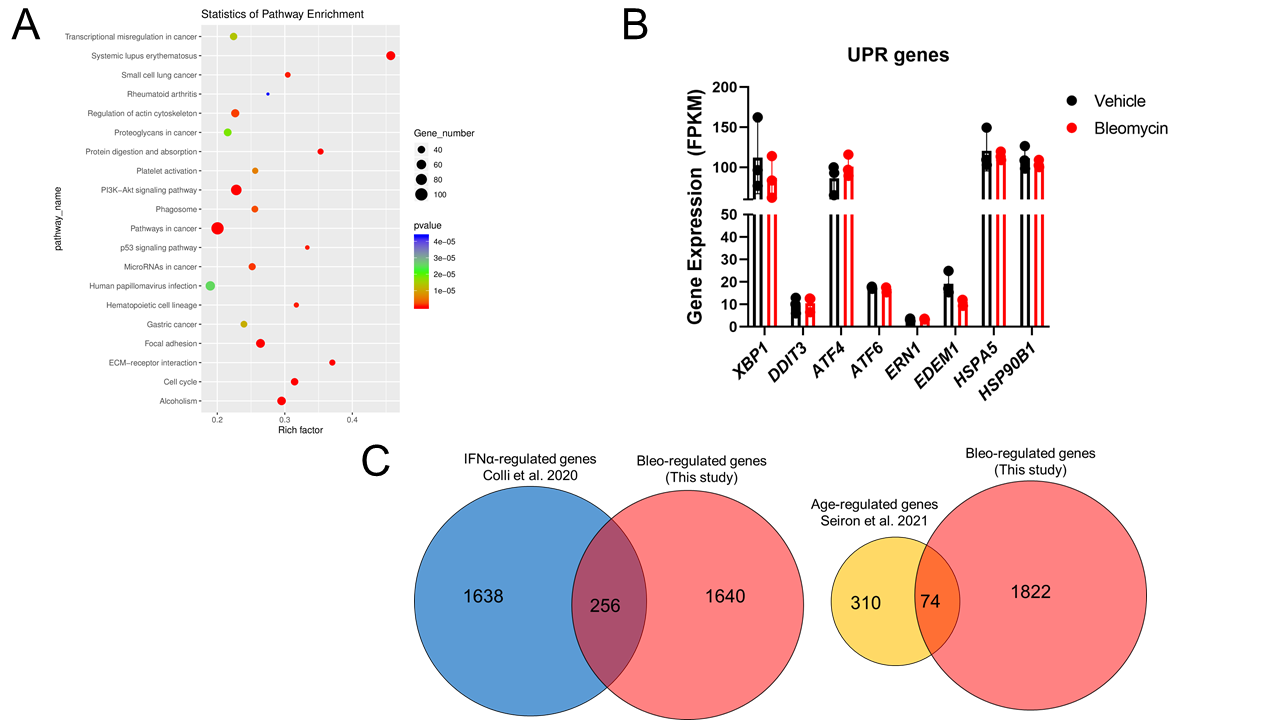

**Supplementary Figure 5. RNA-seq analysis of bleo-treated human islets.** (A) KEGG pathway analysis for significantly enriched terms in differentially expressed genes from bleo-treated versus control human islet RNA-seq as in Figure 4A. Full list of gene expression data is found in Supplementary Table 1. (B) Selected UPR genes from Supplementary Table 1 are plotted. No significant differences in any of these genes were found. (C) List of differentially expressed genes in this RNA-seq dataset (Bleo-regulated genes) was compared with whole islet RNA-seq datasets that identified IFNα-regulated genes (Colli et al. 2020 Nat Comms) or Age-regulated genes (Seiron et al. 2021 PLOS ONE). Genes identified in common are shown in overlapped region of Venn diagrams and listed in Supplementary Table 1.

**Supplementary Tables**

**Supplementary Table 1.** RNA-seq gene expression levels (FPKM) for vehicle and bleomycin treated islets, along with differentially expressed genes in common with natural aging (Seiron et al. 2021 PLOS ONE) and IFNα cytokine exposure (Colli et al. 2020 Nat Comms).

**Supplementary Table 2.** Known and predicted Long noncoding RNAs differentially expressed between vehicle and bleomycin-treated human islets.

**Supplementary Table 3.** Antibodies and qPCR primer sequences used in this study.
